# In Vivo Localization of Deep Brain Implants in Mice

**DOI:** 10.1101/2020.03.28.011858

**Authors:** Bálint Király, Diána Balázsfi, Ildikó Horváth, Nicola Solari, Katalin Sviatkó, Katalin Lengyel, Eszter Birtalan, Domokos Máthé, Krisztián Szigeti, Balázs Hangya

**Affiliations:** Lendület Laboratory of Systems Neuroscience, Institute of Experimental Medicine, Hungarian Academy of Sciences, Budapest, Hungary; Department of Biophysics and Radiation Biology, Semmelweis University, Budapest, Hungary; Department of Biological Physics, Eötvös Loránd University, Budapest, Hungary; János Szentágothai Doctoral School of Neurosciences, Semmelweis University, Budapest, Hungary; CROmed Translational Research Centers, Budapest, Hungary

**Author notes:** Correspondence: *Balázs Hangya,.

## Abstract

Electrophysiology provides a direct readout of neuronal activity at a temporal precision only limited by the sampling rate. However, interrogating deep brain structures, implanting multiple targets or aiming at unusual angles still poses significant challenges even for expert operators, and errors are only discovered by post-hoc histological reconstruction. Here, we propose a method combining the high-resolution information about bone landmarks provided by micro-CT scanning with the soft tissue contrast of the MRI, which allowed us to precisely localize electrodes and optic fibers in mice *in vivo*. This enables arbitrating the success of implantation directly after surgery with a precision comparable to the gold standard histological reconstruction. Adjustment of the recording depth with electrode microdrives or early termination of unsuccessful experiments saves many working hours, while fast 3-dimensional feedback helps surgeons to avoid systematic errors. Increased aiming precision will allow more precise targeting of small or deep brain nuclei and multiple targeting of specific cortical layers.

## Introduction

One of the most common approaches of modern experimental neuroscience is recording or influencing the activity of small brain areas in live laboratory animals. For instance, electrophysiology provides a direct readout of neuronal firing at the single cell level with high temporal resolution^1–3^. Additionally, the recent advent of optogenetics allows activating or suppressing genetically defined groups of neurons via the cell type specific expression of light-gated ion channels and light delivery via optic fibers^4,5^. The combination of the two enables optogenetic tagging, i.e. light-guided assessment of neuron identity during extracellular recordings from awake behaving animals^6–9^. Finally, fiber photometry techniques are on the rise, requiring fiber optics implants similar to optogenetic experiments^10–13^.

Currently, there is strong focus on engineering ever-evolving opsin actuators, optimizing viral vectors for their delivery and improving recording electrodes in terms of channel count and arrangement, tissue damage and durability^5,14–20^. However, experimental success rates are still largely limited by operation and targeting techniques, which remained essentially unchanged since the introduction of stereotaxic surgeries.

Mice represent an attractive model due to their relatively low price, short generation time, genetic tractability, complex behaviors and a level of reported homology between rodent and primate brains. However, surgeries on the mouse brain have gradually become more demanding and error-prone as the field moves towards more complex experiments, aiming at small and often multiple targets simultaneously, hitting the same target multiple times (e.g. virus injection and optic fiber implantation) or implanting at unusual angles to avoid large blood vessel. Targeting deep brain structures poses a significant challenge, as slight deviations from ideal implanting directions or coordinates often leads to missing small deep nuclei by a few hundred microns even in the hands of expert operators. Errors are only discovered at the end of the experiments by post mortem histological reconstruction, often resulting in months of wasted experimental work and lab resources, especially when behavioral training is involved.

In order to improve the efficiency of these experiments, we developed a new procedure in mice inspired by techniques used in human deep brain surgery and brain radiation therapy^21–24^. CT imaging has been used to localize 50 μm diameter electrodes post mortem with an accuracy sufficient for larger (>1000 μm) structures of the rat brain^25^ and large (200 μm diameter) electrodes and lesion sites were localized with CT and MRI in rats *in vivo*^26^. However, the two-fold size difference between mouse and rat brains and the need for targeting small nuclei makes sufficiently detailed imaging and localization challenging and do not allow direct implementation of these methods for precisely localizing small diameter implants in the mouse brain. To overcome these limitation, we used high-resolution (19 μm) micro-CT imaging that allowed accurate measurements of tetrode electrodes (12.7 μm single wire diameter), silicon probes (Buzsáki-probe with a 52 × 15 μm shank) and optical fibers (250 μm diameter) with respect to bone landmarks. However, CT does not provide soft tissue contrast, making it difficult to judge whether the implant is in the target area. Therefore, we performed structural MRI scans that provide excellent soft tissue contrast and merged them with the micro-CT images, similar to human surgery techniques. The procedure consists of pre-operative and post-operative imaging, which is followed by the registration (i.e. aligning) of the images with an atlas coordinate system (Fig. 1), providing localization of implants with a precision that matches post hoc histological reconstruction. This non-invasive *in vivo* localization technique enables verifying the success of implantation directly after surgery and increases the efficiency of the experiment by (i) adjusting the recording depth to reach target, (ii) terminate in case of mistargeting or other problems and (iii) systematically improve surgery skills by providing fast feedback. We estimate that this method may save up to an order of 1000 working hours per chronic recording/optogenetics/fiber photometry projects, depending on the difficulty of the surgery and the length of behavior training or other repeated procedures.

**Figure 1.**
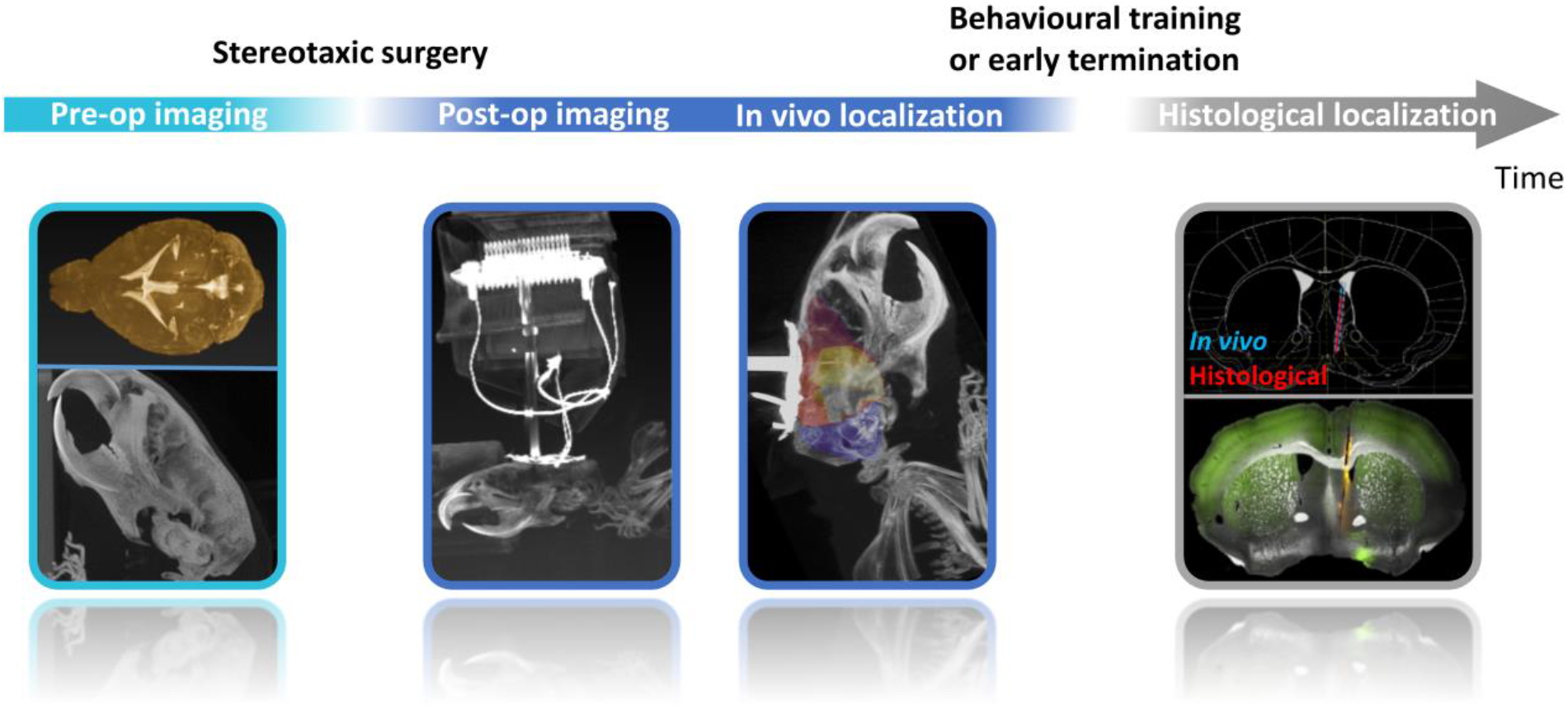
*In vivo* localization workflow. A few days before the surgery, pre-operative CT and MRI scanning is performed providing anatomical information about bone landmarks and the brain. After 4-12 days recovery post the stereotaxic surgery, post-operative CT images are taken and the position of the implant is determined in a co-registered atlas coordinate system. Based on the results, the trajectory of the implant is extrapolated, depth of the implant may be adjusted, or in case of mistargeting, early termination is considered. We verified the accuracy of the *in vivo* localization using post hoc histology.

## Results

### CT-based implant localization

High-resolution CT images of the skull allow image registration to a common stereotaxic coordinate system based on bone landmarks. As important bone landmarks are often masked by surgically applied radiopaque dental adhesives, these scans are ideally performed in intact animals. Therefore, we conducted pre-operative micro-CT measurements at 35 μm resolution of anesthetized mice fixed by the top incisor teeth in a specialized isoflurane mask to avoid small movements and displacements (Fig. 2a). Next, we aligned the imaging planes with canonical 3D axes used in brain atlas coordinate systems, similar to stereotactic adjustment of skull position, often referred to as ‘leveling’ (Fig. 2b). Leveling of the transversal plane was based on Bregma and Lambda points on the surface of the calvaria (blue dashed lines in Fig. 2b). Bregma and Lambda (blue and yellow arrows in Fig. 2b) were defined as the midpoints of the best fit curves on the coronal and the lambdoid sutures, respectively^27^. We used a collection of symmetrical bone structures to aid leveling of the sagittal and coronal planes including the squamous, petrous and tympanic part of temporal bone, the zygomatic process, the temporal line, the basisphenoid bone, the presphenoid bone, the tympanic bulla, the ear canal, inner ear structures and other symmetrical points of the cranium (yellow dashed lines in Fig. 2b).

**Figure 2.**
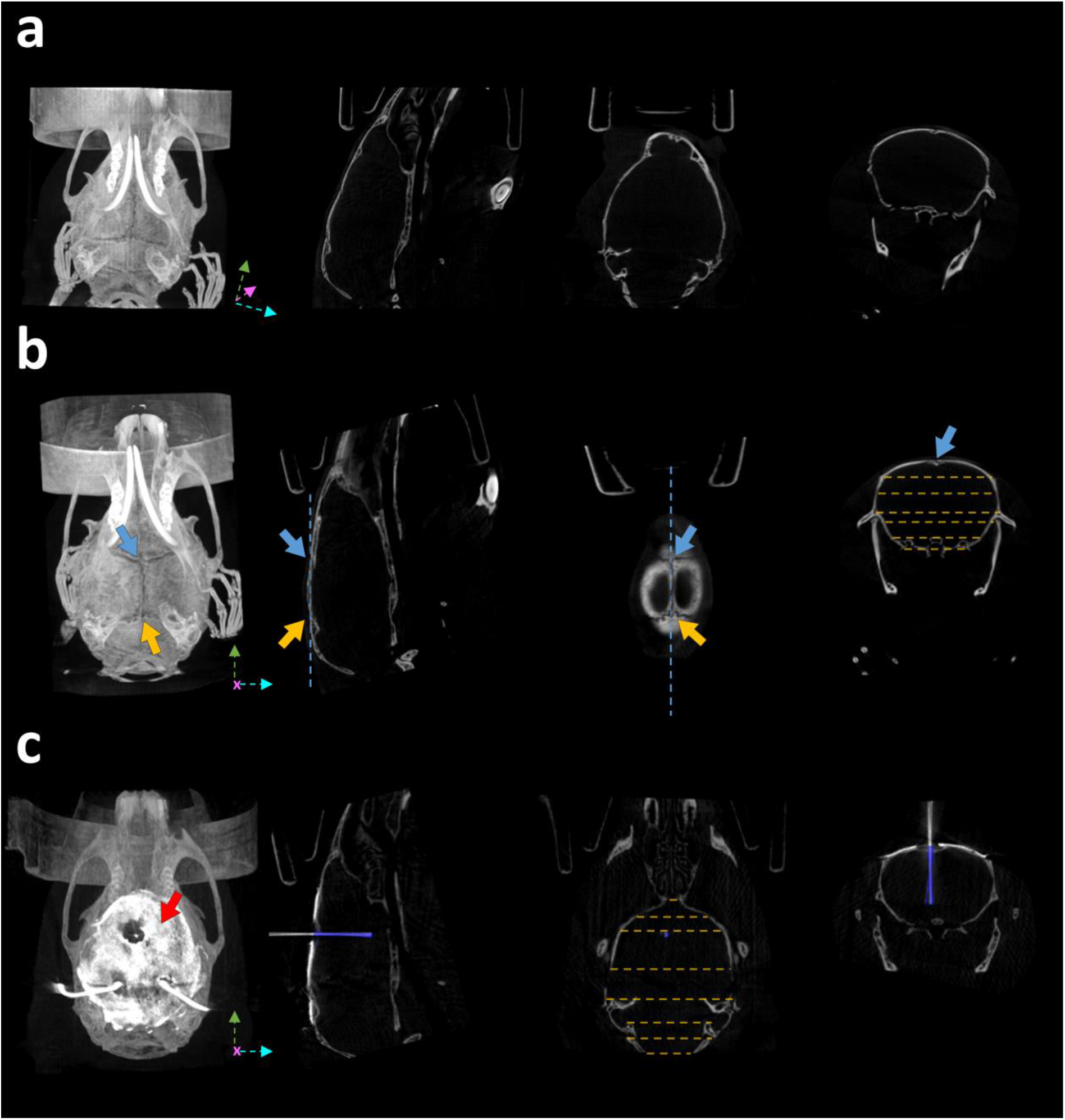
CT-based implant localization. **a** Raw pre-operative CT image. Canonical 3D axes used in brain atlas coordinate systems (green, pink and cyan arrow) are not aligned with the coordinate system of the CT image. **b** The pre-operative CT image aligned with the atlas coordinate system, after leveling the Bregma (blue arrows) with the Lambda (yellow arrows) in the sagittal plane (blue dashed lines) and symmetrical bone structures in the coronal and horizontal planes (yellow dashed lines). **c** Post-operative CT image after the implantation of an 8-tetrode microdrive, co-registered with the pre-operative CT image. The skull was covered with a radiopaque dental adhesive (red arrow), which hides Bregma and Lambda. The implant was segmented (blue area) with an intensity threshold applied on a whole brain ROI. Images in each row from left to right, maximum intensity projection, sagittal slice, horizontal slice, coronal slice.

Standard stereotactic surgeries were performed by experienced operators to implant chronic electrode drives housing 8 tetrodes and an optic fiber. During surgery, either the horizontal limb of the diagonal band of Broca (HDB, n = 5), a deep brain target in the basal forebrain area (+0.74 mm antero-posterior, 0.6 mm lateral and 5.0 mm dorso-ventral from Bregma; see Methods), or the medial septum of the basal forebrain (MS, n = 2) was targeted (+0.9 mm antero-posterior, 0.1 mm lateral and 3.9 mm dorso-ventral from Bregma). MS is a midline structure and thus it was implanted at a 14 degrees angle with respect to the vertical axis in the coronal plane to avoid the sinus sagittalis superior.

To accurately localize the implanted electrodes, we performed post-operative CT scanning at 19 μm resolution after a 4-12 days recovery period using the same procedure as detailed above (Fig. 2c). To minimize shadow artifacts introduced by metal objects in the region of interest of the CT images, we positioned the metal parts of the drive, i.e. wiring and pins of the electrode interface board and occasional hypodermic tubing or head-bars, to point away from the approximately coronal plane in which the X-ray source was rotating. Next, the post-operative CT scan was co-registered with the pre-operative one that had already been aligned to the stereotaxic coordinate system by matching corresponding bone structures using rotation and translation operations on the post-operative image (Fig. 2c). Finally, the implant was segmented with an intensity threshold applied on a whole brain region of interest (ROI) (dark blue area in Fig. 2c). As a result, we were able to read the stereotaxic coordinates of the tip of the implant relative to the Bregma with reference to brain atlas coordinates (e.g. Paxinos and Franklin’s the Mouse Brain in Stereotaxic Coordinates^27^) and calculate the angle of the implant trajectory. This also allowed us to use trigonometric functions to calculate the future position of the implant along its projected trajectory.

### Implant localization based on CT-MRI fusion

Previously, we registered the tips of the implanted electrodes to stereotaxic atlas coordinates based on the combination of pre- and post-operative CT scans. However, as CT provides practically no soft tissue contrast in the mouse brain, it is hard to judge the area localization of the electrode as well as the distance from neighboring structures in three dimensions based on the images. Additionally, CT-based registration relies on a few bone landmarks without any possibility of correction based on brain structures. Consequently, the above method cannot take individual variability of brain morphology into account, and strongly depends on the somewhat subjective identification of the Bregma point.

Therefore, we further developed our localization method based on the CT-MRI fusion technique routinely applied during human Deep Brain Stimulation (DBS) surgeries and brain radiation therapy ^21–24^. In addition to the pre-operative CT, we performed a T1-weighted pre-operative MRI scan directly after the CT-scan. To aid the subsequent CT-MRI fusion, mice were kept in the same scanning bed across the two modalities, avoiding postural changes and large disparities between the imaging planes. Next, we developed a multi-step procedure to co-register our scans with a 3D brain atlas^28^, enabled by the soft tissue information obtained from the pre-operative MRI (Fig. 3). As an important rule to guide precise image registration, in each of the following steps, either images from the same modality or images from the same animal acquired in the exact same position were co-registered.

i. First, the pre-operative CT image was aligned to stereotaxic axes as described in the previous section.
ii. Second, the pre-operative CT and MRI images were co-registered (Fig. 3a). This could be achieved in a relatively simple fusion procedure involving small translations applied on the MRI image, since the two scans were acquired in the same scanning bed while keeping the position of the head of the animal fixed. The transformations were based on the skull, which provides high-contrast signal in CT images while appears as a distinct lack of signal between soft tissue structures in the MRI.
iii. Third, the pre-operative MRI image was co-registered with the qualitatively best matching sample MRI image of the 3D atlas, based on the contour of the brain and the position of the ventricles in multiple planes (Fig. 3a, blue arrows). This was achieved by a combination of translations and affine transformations, the latter of which ranged up to a few percent. Note that this was the only step in the protocol that required a non-Euclidean transformation. If entirely uniform fitting was not achievable across the whole brain, we paid special attention to the perfect alignment of the structures close to the area of interest, i.e. the surgical target area.
iv. Fourth, we applied the transformation matrix that realized the best alignment in step (iii) on the MRI atlas, thus obtaining registration of individual mouse MRI scans to a reference MRI-atlas (Fig. 3b). Thus, the first four steps result in the co-registration of the pre-operative CT and the MRI-atlas, providing individualized brain area information in a stereotaxic coordinate frame aligned to bone landmarks.
v. Fifth, we verified the fitting of characteristic bone landmarks such as the Lambda and the sinus frontalis (Fig. 3c, yellow arrows) with the contour of the atlas, and in case of misalignment, the first four steps were revised.
vi. Sixth, the post-operative CT image was co-registered with the pre-operative CT as described in the previous section, resulting in a fusion between the post-operative CT and the reference MRI-atlas (Fig. 3c). In the coronal slices containing the tip of the implant, the corresponding coronal plane of the anatomically more detailed and readily scalable Paxinos atlas was fitted on the three-dimensional atlas (Fig. 3d), which aided further quantitative comparisons (see below).

**Figure 3.**
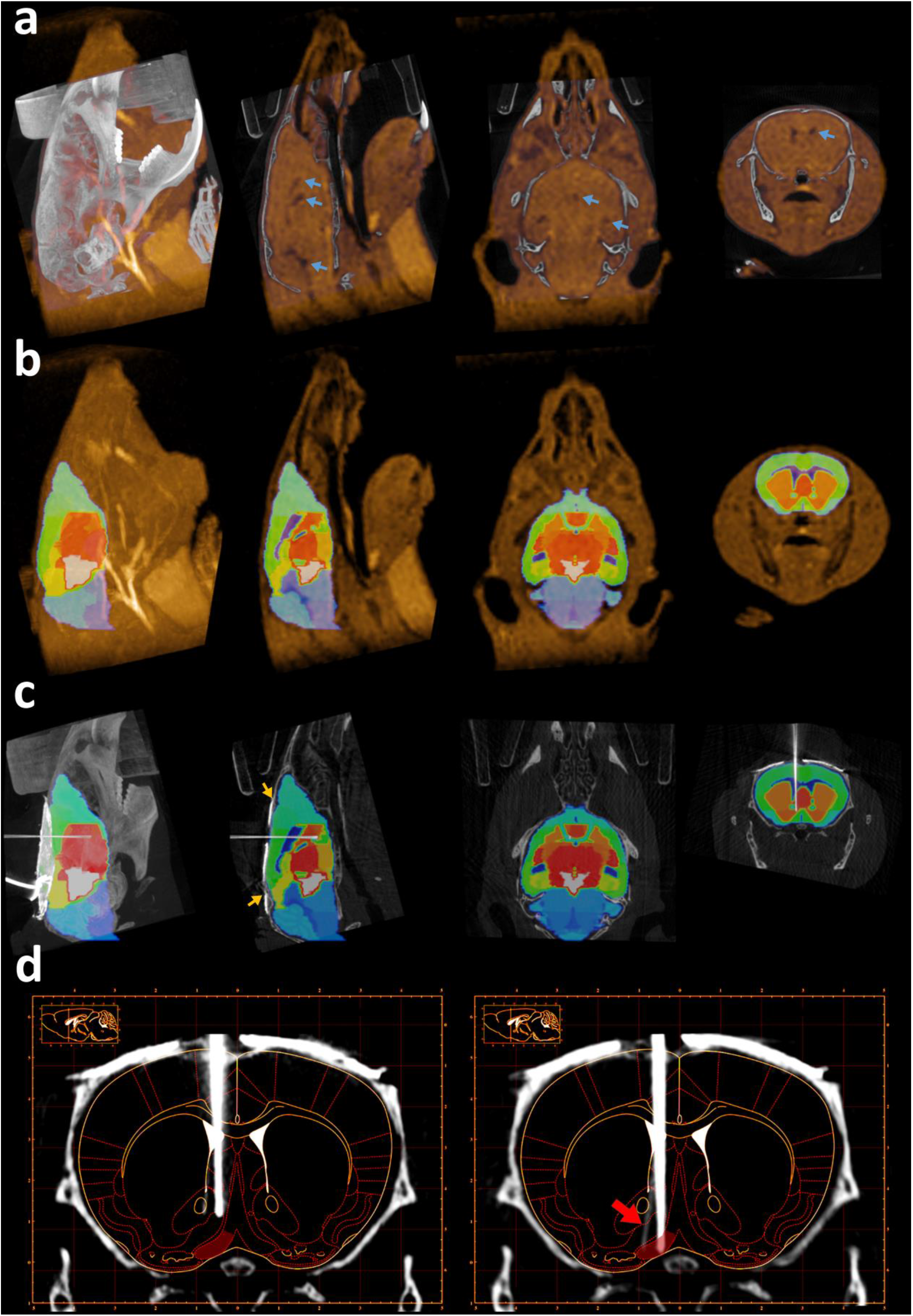
Implant localization based on CT-MRI fusion. **a** Fused pre-operative CT and MRI images (same mouse as in Fig. 2). Blue arrows show the ventricles on the MRI image used for atlas co-registration. Images from left to right, maximum intensity projection, sagittal slice, horizontal slice, coronal slice. **b** The 3D atlas co-registered with the MRI image. **c** The 3D atlas co-registered with the post-operative CT image. Yellow arrows show Lambda and sinus frontalis. **d** The corresponding coronal plane of the Paxinos atlas fitted on a coronal slice containing the tip of the implant after surgery (left) and at the end of the electrophysiology experiment (right). The figures clearly show that the implanted tetrodes went through the target area (HDB, red shading) during the experiment. The red arrow shows a tetrode that separated from the bundle.

This procedure resulted in a three-dimensional image displaying the trajectory of the implant in the brain. Similar to the planning phase of human DBS surgeries, we could determine the exact position of the implant relative to the target area and predict the future positions along the planned descent. This allowed more precise knowledge of the recording position throughout the entire experiment, and – in case of a mistargeting – provided a chance to determine the exact source of error (see below).

### Localization of multiple-targeting implants, silicon probes and optic fibers

We have demonstrated precise, non-invasive *in vivo* localization of implanted tetrode bundles targeting single deep brain regions using structural imaging. As questions about information transfer and coordinated processing across distant brain areas are increasingly coming into the focus of research, targeting multiple brain areas simultaneously is becoming common practice. Therefore, we tested whether our previously demonstrated localization technique could be applied to implants with multiple targets. To this end, we implanted a custom-built dual electrode drive housing 8-8 tetrodes with 1-1 optic fiber at a fixed distance of 3.85 mm for simultaneous targeting of two subcortical neuromodulatory centers, the HDB of the basal forebrain and the midbrain ventral tegmental area (VTA; −3.1 mm antero-posterior, 0.6 mm lateral and 4.05 mm dorso-ventral from Bregma). The presence of more metal parts in these drives introduced somewhat more noise in the post-operative CT images compared to single tetrode bundle implants. Nevertheless, we were able to reconstruct both tetrode trajectories and localize the tips by the *in vivo* technique (Fig. 4a).

**Figure 4.**
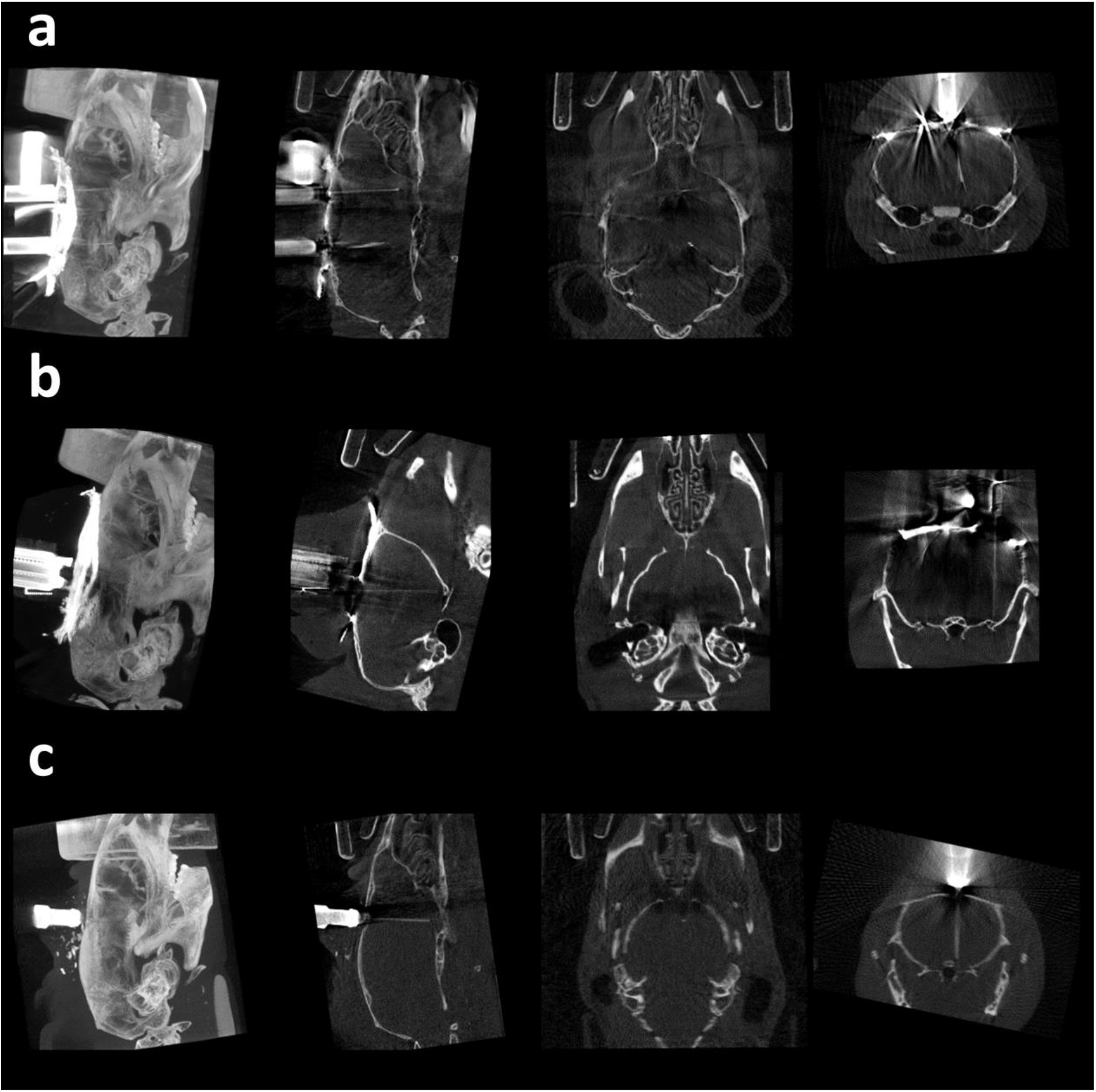
Localization of multiple-targeting implants, silicon probes and optic fibers. **a** CT image of a dual tetrode drive implanted in HDB and VTA. Images from left to right, maximum intensity projection, sagittal slice, horizontal slice, coronal slice. **b** CT image of a single shank silicon probe implanted in the ventral CA3. **c** CT image of a non-metalic optic fiber implanted in HDB.

While tetrode recordings may be considered the mainstay of *in vivo* extracellular electrophysiology, a large variety of silicon probe electrode designs are gaining popularity due to their high channel count, customized contact site configuration and ease of use. Therefore, we tested the *in vivo* localization technique on a silicone probe implant as well. A Buzsaki-type silicone probe (52 μm width and 15 μm thickness, one of the thinnest commercially available) was lowered into the ventral CA3 area of the hippocampus (−2.5 mm antero-posterior, 2 mm lateral and 4.7 mm dorso-ventral from Bregma). The silicone probe trajectory and tip were successfully visualized by micro-CT imaging, showing that the localization technique is readily applicable to silicone probe electrodes (Fig. 4b).

We have demonstrated precise localization of metal electrodes using their high contrast signal in the CT images. With the rapid progress of optical methods for imaging and manipulating neural activity including optogenetics and fiber photometry, optic fiber implants have become common in chronic *in vivo* experiments. Based on the approximate Hounsfield unit (HU) of different glass and plastic objects 29,30, silica glass and plastic optic fibers are expected to induce a signal intensity between the ones induced by bone structures and soft tissue. We implanted (n = 2) mice with a 250 μm core multimode silica optic fiber targeting the HDB or the VTA, suitable for stimulating or inhibiting neural activity in optogenetic experiments, and subsequently tested the localization method. While the non-metallic optic fiber indeed provided a lower-contrast signal compared to the metal electrodes, micro-CT imaging proved to be sufficient to reveal the position of single optic fibers on the soft tissue background (Fig. 4c).

### Quantification of localization accuracy using gold-standard histology

Post mortem histology and anatomical reconstruction of the implant’s track is considered as gold standard technique for implant localization. We tested the accuracy of *in vivo* localization against this gold standard by examining the outcome of 7 tetrode and 1 dual tetrode surgeries targeting deep brain neuromodulatory nuclei such as the horizontal limb of the diagonal band of Broca (HDB, n = 6), the medial septum (MS, n = 2) and the ventral tegmental area (VTA, n = 1).

After the *in vivo* experiments, mice were anesthetized and an electrolytic lesion procedure was applied to mark the location of the electrode tips, then mice were perfused transcardially and 50 μm coronal sections were prepared using standard histology techniques (see Methods). Implant tracks were examined in bright and dark field microscopy images based on the electrolytic lesion site and the trace of the implant along its decent. Tracks were also visualized in fluoromicrographs by red fluorescent DiI applied on the implants. Atlas sections^27^ at corresponding coronal levels were aligned with the microscopy images using linear scaling and rotation, which approach provided atlas coordinates of the tip and track of the implants (Fig. 5a). Histological reconstruction was then compared with the *in vivo* localization (Fig. 5b and 5c).

**Figure 5.**
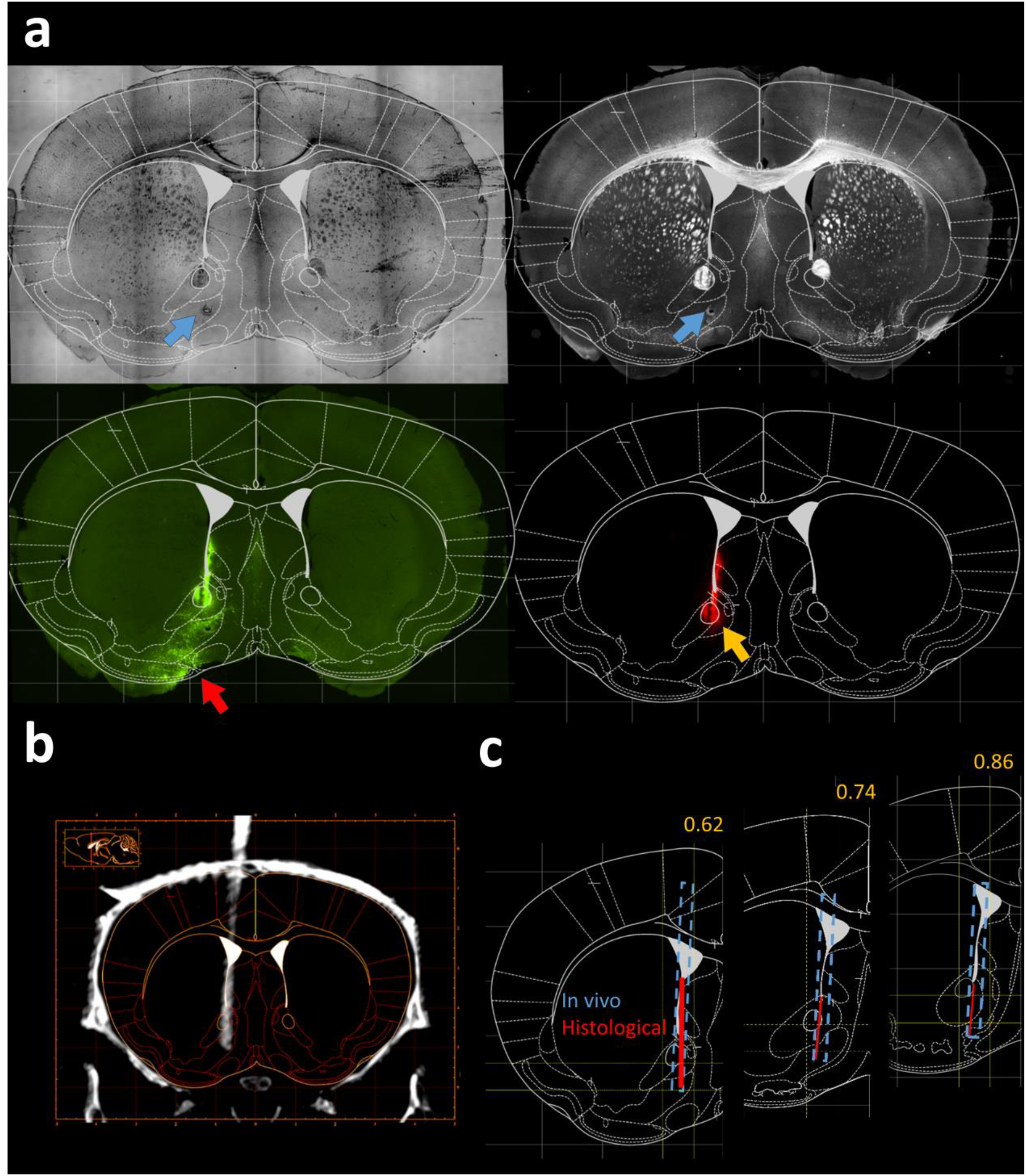
Track reconstruction with post hoc histology in comparison with in vivo localization. **a** The Paxinos atlas was fitted on coronal sections based on anatomical landmarks identified in the bright-field (top left) and dark-filed (top right) images. The atlas image was overlaid on fluorescent images showing the virus infection (eYFP) at the target area (red arrow, bottom left) and the implant track (DiI, yellow arrow, bottom right). The tip of the electrode was marked with an additional electrolytic lesion (blue arrows). **b** Corresponding coronal CT slice with the implant trajectory localized *in vivo*. **c** Comparison of the histologically reconstructed implant track (red line) and the *in vivo* localization (blue dashed line). Yellow numbers refer to the antero-posterior coordinates.

To go beyond this qualitative comparison and provide comparable measures of targeting accuracy, we quantified the offset relative to target along the three cardinal axes both with *in vivo* localization and histological reconstruction. The location of the implant tip was assessed as a range of coordinates in the antero-posterior and medio-lateral directions spanned by the full extent of the radiodense area (CT) or the track in the tissue (histology), and characterized by a single dorso-ventral coordinate corresponding to the most ventral point (Fig. 6a and Supplementary Fig. 1a). As expected, offset measures were strongly correlated between the *in vivo* and the histological localization in all dimensions (Pearson correlation coefficient r > 0.9, Supplementary Fig. 2a).

**Figure 6.**
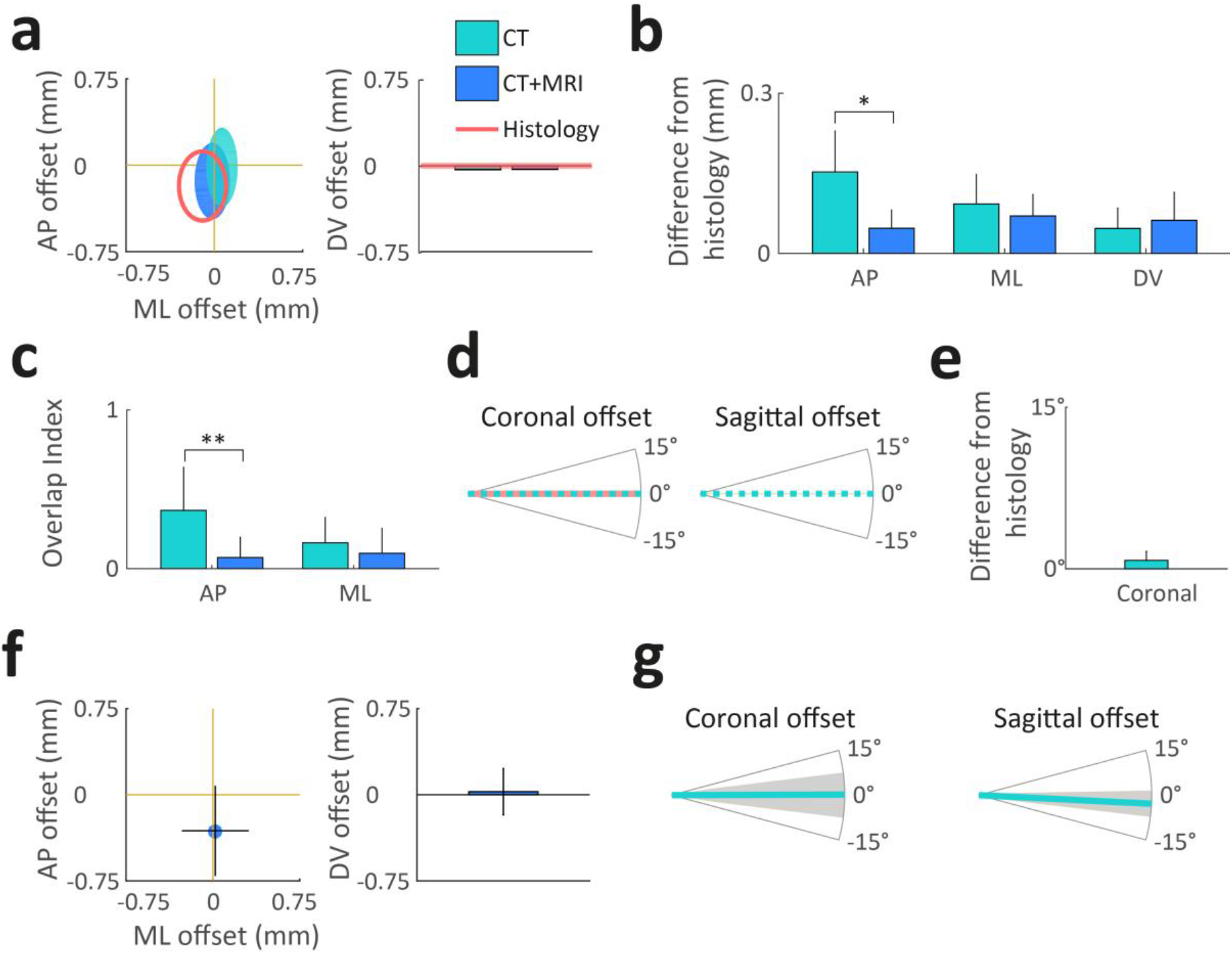
Quantification of localization accuracy. **a** Antero-posterior (AP), medio-lateral (ML) and dorso-ventral (DV) offset from the target coordinate for an example animal. Teal, CT-based coordinate system alignment; blue, CT-MRI fusion-based atlas co-registration; salmon, histological track reconstruction. Ellipses on the left represent the area covered by the tip of the implant in the horizontal plane. **b** Mean absolute difference of tip coordinates in three directions with respect to histology. CT-MRI fusion-based localization was significantly closer to histology results in AP direction. *, p<0.005; two-tailed Wilcoxon signed rank test. **c** Distance from histology based on the overlap of implanted area. CT-MRI fusion-based localization was significantly closer to histology results in the AP direction. **, p<0.01; two-tailed Wilcoxon signed rank test. **d** Coronal and sagittal offset from the planned direction of trajectory for the same example animal as in panel a. Implant directions were quantified based on the CT images. Histology only provides coronal offset measures. **e** Mean absolute difference of CT-based coronal angle compared to histology. **f** Mean offset from target of the centroid of the localized areas with CT-MRI fusion-based localization. **g** Mean offset from the coronal and sagittal target direction with CT-based localization. Anterior, lateral and dorsal directions were defined as the positive directions. Error bars and gray shading show standard deviation.

To quantify the accuracy of *in vivo* localization, we calculated the mean absolute difference between *in vivo* and histology coordinates (Fig. 6b). In the dorso-ventral and medio-lateral directions, *in vivo* localization, either based on CT measurement only or relying on the CT-MRI fusion, showed an average deviation from histology of less than 0.1 mm (maximum, 0.17 with CT and 0.15 with CT-MRI fusion; no significant difference between methods or directions, p > 0.1 two-tailed Wilcoxon signed rank test). In the antero-posterior direction, the CT-MRI fusion technique proved to be significantly closer to the histological reconstruction (average, 0.05; maximum, 0.09) than the CT-based coordinate system alignment (average, 0.15; maximum, 0.29; two-tailed Wilcoxon signed rank test, p<0.005).

Next, we compared the area overlap between *in vivo* and histological localization in the medio-lateral and antero-posterior directions, quantified as 1 minus the length of the intersecting range normalized by the length of the shorter range (Overlap Index; Fig. 6c). Similar to the mean difference comparison in Fig. 6b, the Overlap Index showed significantly better localization with CT-MRI fusion compared to CT alone in the antero-posterior direction (p < 0.01), while there was no significant difference in the medio-lateral direction (p > 0.1, two-tailed Wilcoxon signed rank test).

Finally, we compared the angle of the implant trajectory. The *in vivo* localization method provided quantitative information on the direction of the trajectory in both the coronal and sagittal planes based on the CT scans, while the histological reconstruction only provided angle measurements in the coronal plane (Fig. 6d and Supplementary Fig. 1b). Coronal offset from the target direction was strongly correlated between the *in vivo* and the histological localization (Pearson correlation coefficient r > 0.9, Supplementary Fig. 2b) and we only detected negligible mean difference in the offsets (average, 0.78°; maximum, 2°; no significant difference from 0°, p > 0.1, t-test; Fig. 6e).

These results suggest that the *in vivo* localization method based on CT-MRI fusion has a comparable accuracy with the gold standard histological track reconstruction. While localization based on CT only provided comparable results in the medio-lateral and dorso-ventral directions, it was significantly inferior to the fusion approach in the antero-posterior direction. The source of this difference might be that (i) Bregma was not unambiguously determined by the sutures or (ii) the size of the brain was different from the size defined by the Paxinos atlas in the antero-posterior direction.

### Modes of failure revealed by the in vivo localization

Next, we used *in vivo* localization to identify the exact source of targeting error that typically remains hidden when using histology. Quantification of implantation tracks showed that our surgeons tended to implant more posterior than planned (−0.31 ± 0.39mm, mean ± standard deviation, significantly more posterior then the target, one-tailed t-test, p < 0.05; Fig. 6f). This was due to a slight angle in the parasagittal plane (−2.84° ± 3.86°, significantly more posterior than the dorso-ventral direction, one-tailed t-test, p < 0.05; Fig. 6g) that positively correlated with the targeting offset (Pearson correlation coefficient r > 0.75, Supplementary Fig. 2c; the bent implant was excluded, see below; mean direction used for the dual implant). These results suggest a systematical error in the stereotaxic leveling of the skull during surgery, probably owing to the ambiguity of the Lambda point that has a strong influence on antero-posterior leveling. There was no systematic deviation from the medio-lateral target coordinate and from the preferred direction in the coronal plane (0.13°, Fig. 6d), although the standard deviation was relatively large (± 6.73°). As expected, the coronal angle offset was positively correlated with the medio-lateral offset of the tip (Pearson correlation coefficient r > 0.78, Supplementary Fig. 2c).

Sporadic errors also contributed to the relatively large standard deviations of trajectory angles. In one case, the direction of implantation was more than 2° anterior; the *in vivo* imaging revealed that this was due to bending of the tetrode bundle during the implantation procedure (Fig. 7a, blue arrow). In some surgeries the tetrode bundle was not parallel to the guiding cannula (Fig. 7b, red dashed line), suggesting errors in the drive assembly process. We also found a damaged implant with a kink in the tetrode bundle outside the brain (Fig. 7c, yellow arrow), which either suggests a problem with the micro-drive building procedure or failing to prevent dental cement from reaching the craniotomy. In a further case, the implanted electrodes covered an unusually large area with individual tetrodes and the optic fiber spanning multiple brain regions (Fig. 7d, blue dashed circle), indicating a larger than optimal collateral spread of the tetrode bundle.

**Figure 7.**
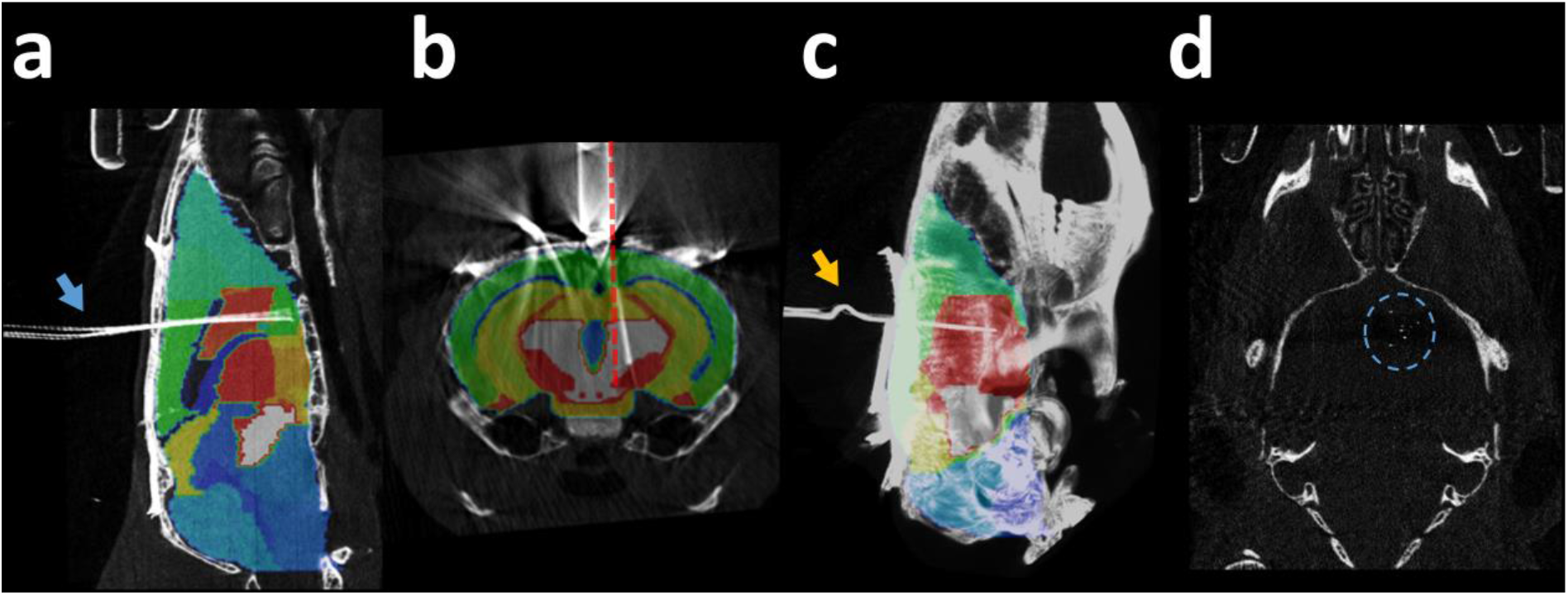
Modes of failure revealed by the *in vivo* localization. **a** Bending of the tetrode bundle (blue arrow). **b** Tetrode bundle not parallel with the cannula (red dashed line, the cannula was aligned with the dorso-ventral direction during the stereotaxic surgery). **c** Kink on the tetrodes outside the brain (yellow arrow). **d** Over-dispersion of the tetrodes, covering multiple adjacent brain areas (blue dashed circle).

### Optimization of image quality and radiation dose

In micro-CT imaging, high resolution is achieved by lowering the distance between the source and the object, which is inversely proportional to square of radiation dose. In addition, a better signal to noise ratio (SNR) can be achieved by longer exposition time and an increased number of projections. Therefore, micro-CT imaging at high resolution and SNR may involve relatively high radiation doses, thus possible adverse effects of radiation cannot fully be ruled out. As this issue has not hitherto been investigated in depth, we performed dose measurements and health monitoring at a wide range of parameters to optimize micro-CT imaging with respect to this image quality - radiation dose trade-off.

We tested several acquisition settings for magnification, number of projections and exposure time (n = 3 mice for each setting; Fig. 8a). Parameters that did not affect the information content of the acquired images significantly, e.g. photon energy in the available 45 kVp-65 kVp range, were set to minimize the radiation dose. The tested settings are referred to by numbers (i) to (v) as shown in Fig. 8a. Radiation dose was measured with termoluminescent detectors (TLD) attached to the neck of the animals as close to the skull as possible, where the highest dose was expected based on spatial considerations. The measured dose of a single scan ranged from 4078 to 79 mGy across settings (i) to (v). All settings were found to be suitable for localizing all relevant bone structures. Settings (i), (ii) and (iii) visualized the position of tetrode implants with the precision required for the localization methods. For localizing the silicone probe, the better SNR of settings (i) and (ii) were necessary, while the bigger diameter optic fiber implant could be reliably localized with the lower resolution of setting (iv), too (Supplementary Fig. 3). Most settings did not produce observable adverse health effects during the four weeks of follow-up period of the animals. However, with setting (i) representing the highest radiation dose, we observed a slight loss of hair and skin irritation around the neck of the animals after 2-3 weeks. Therefore, we restricted further scans to settings (ii) to (v).

**Figure 8.**
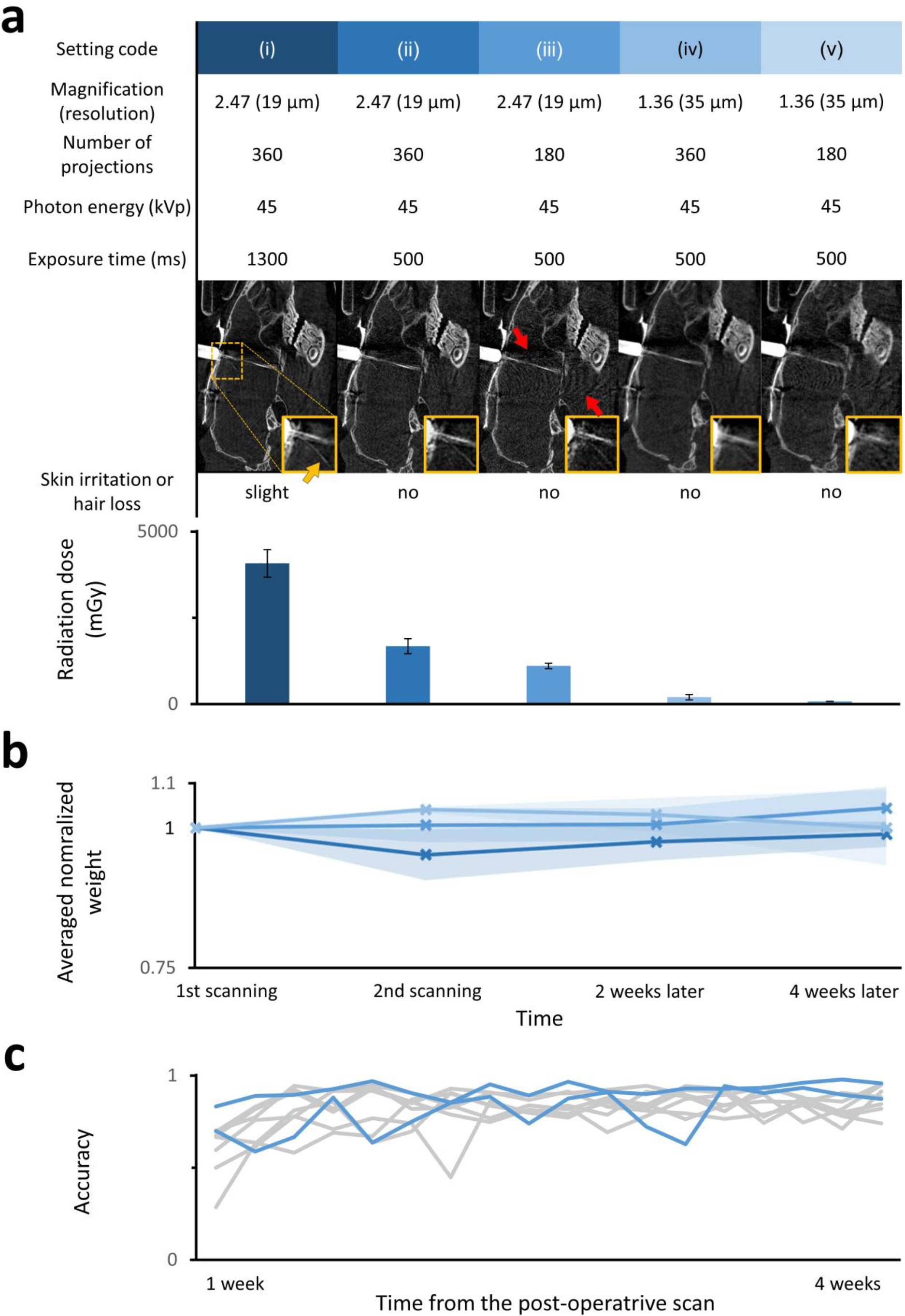
Optimization of image quality and radiation dose. **a** Top, acquisition settings for CT; middle, corresponding image quality of a tetrode drive implant (for slilicon probe and optical fiber implants see also Supplementary Fig. 3); bottom, radiation effects and measured doses (n = 3 mice per group). The yellow arrow shows a bent single tetrode on the magnified image, indicating the spatial resolution of the different images. Red arrows show artefacts caused by the metal parts of the microdrive, amplified if the exposure time and the number of projections are decreased. Error bars show standard deviation. **b** Average normalized body weight of mice (n=3 per group) exposed to two CT scans separated by a one-week pause, simulating the protocol used for the localization procedure. The first scans were performed with setting (v) and the color code of panel a indicates settings for the second scan. Error shades show standard deviation. **c** Performance in a standard 5-choice serial reaction time task of scanned animals (n = 2, settings (v) and (iii), blue) did not differ from that of the control population (n = 8, gray).

Next, we subjected another cohort of animals (n = 3 mice for each group) to the full protocol of two micro-CT scans separated by one week and monitored the weight and general condition of the animals. In all groups, the first scan was performed with setting (v) characterized by the lowest, negligible dose that was still suitable for localizing the relevant bone structures. The second scans were performed with settings (ii) to (iv). None of the animals showed any signs of skin irritation, hair loss or drop in weight (Fig. 8b). Based on these findings, we suggest that post-operative CT parameters should be adjusted to match the size and radiodensity of the implant, with the lowest possible radiation dose.

Finally, we compared the performance of mice (n = 2) scanned with setting (v) before and setting (iii) after surgery with a control population (n = 8) not exposed to radiation in a standard 5-choice serial reaction time task^31^. Before the surgery, the animals were pre-trained for 5-10 days in an automated system (see Methods). By the end of the pre-training period, all animals learned the task and performed around 80% accuracy. One week after the surgery, post-operative scanning was performed and the training was resumed. Those mice that had been subjected to the scanning procedure performed at a level of accuracy indistinguishable from the control population (Fig. 8c).

## Discussion

Good temporal resolution and direct connection with neural activity make electrophysiology and optogenetics powerful techniques to study or manipulate neural activity of awake behaving animals. Developed recently, fiber photometry experiments provide a cell type specific readout of the ongoing activity of neural populations. When complicated implants and difficult surgeries, often followed by complex and lengthy behavioral training are increasingly commonplace, variability in targeting accuracy of stereotaxic surgeries is becoming rate limiting. Consequently, the lack of *in vivo* methods for localizing implanted electrodes or optic fibers eventually thwarts the effectiveness of these studies.

Structural imaging has been used to localize implants in humans, monkeys and rats. For instance, Borg and colleagues^25^ used micro-CT imaging in rats for localizing 50 μm diameter microwire electrode arrays post mortem with an accuracy sufficient for the larger structures of the rat brain (with at least 1000 μm width and height). Also in rats, Rangarajan et al.^26^ localized 200 μm diameter electrodes and lesions *in vivo* with CT and MRI, respectively. However, *in vivo* localization of thin electrodes or optic fibers implanted to small deep nuclei of the mouse brain still remained a challenge and therefore *in vivo* implant localization has not been performed in mice.

To help resolve this issue, we developed a noninvasive *in vivo* localization technique for radiodense implants in mice by combining different structural imaging modalities. This method largely relies on transforming the co-registered pre- and post-operative micro-CT images to the stereotaxic reference system, capable of determining the coordinates of the implant relative to Bregma (Fig. 2). This method was aided by a pre-operative MRI image, which was co-registered both with a three-dimensional MRI-based atlas and with the post-operative CT image in a multi-step protocol based on anatomical landmarks (Fig. 3). The added benefit of the structural MRI was that it took individual anatomical variation into account and, by introducing soft tissue contrast absent in the CT modality, provided visual three-dimensional information about the trajectory of the implant in the brain. While the CT-based alignment was sufficiently precise to localize small diameter implants in deep brain nuclei with high accuracy, localization based on the CT-MRI fusion was significantly more accurate along the antero-posterior axis in the sagittal plane. Although we did not find a significant difference between the CT- and fusion-based methods in the medio-lateral direction, we note that implants far from the midline could possibly reveal a larger difference.

Verifying the success of implantation directly after surgery using *in vivo*, non-invasive techniques allows precise adjustment of the recording depth with microdrives, increasing targeting precision and thus experiment efficiency. Moreover, if the implant proved to have missed the target location, which may occur in up to 30-60% of surgeries for deep brain targets according to previous^26,32^ and our estimates, early termination saves valuable time, human and other lab resources. For instance, in a previous study^33^ we implanted 32 mice to target the basal forebrain for optogenetic tagging of central cholinergic neurons. Optogenetic effects detected in 26/32 mice indicated that the electrodes and optic fiber approached cholinergic areas, prompting us to perform full histological reconstruction. Post hoc localization determined that the basal forebrain was hit in 19/32 animals. In the remaining 13 mice, 374 recording sessions were performed, each lasting approximately 2 hours that included behavioral training and two optogenetic tagging sessions. Thus, we estimate that early termination based on *in vivo* localization would have saved about 750 net working hours throughout the course of the project, while shortening decent time in the successfully implanted mice could have further optimized experimental time.

We showed that *in vivo* localization based on the CT-MRI fusion technique provides the same level of accuracy as histological track reconstruction (Fig. 6) with comparable time investment (approximately 3 hours per animal). In addition, the *in vivo* method is less subjective and more reliable in the sense that it is free from all the irreversible information loss that may occur during slice preparation, e.g. tissue shrinkage during perfusion, damage caused by removing the implant, suboptimal angle of slice preparation and tissue damage or loss during sectioning, which can be especially problematic when multiple implants are positioned close to each other. We suggest that in projects using *in vivo* localization, it becomes unnecessary to sacrifice the animal immediately after the experiment. This opens the possibility of designing long-term experiments, where the animal may be followed up after the training process, tested for long-term memory or trained on different tasks.

Targeting accuracy depends on a number of factors including scaling errors when using animals of different age, size or strain, errors in aligning to the axis of the atlas system or subtle displacements of the skull due to drilling. Another benefit of visual feedback on surgical procedures is the chance of identifying potential causes of failure including systematic errors committed by the operators that often remain hidden when using histology. Indeed, we discovered a tendency of implanting posterior from the target position due to deviation from vertical trajectory in the sagittal plane that suggested a slight error in bregma-lambda leveling. Other causes of mistargeting included bending or over-spreading of the tetrode bundle and damaged implants that suggested problems with micro-drive building or surgical procedures (Fig. 7).

We showed that the structural information provided by the MRI imaging improved the accuracy of the *in vivo* localization along the antero-posterior axis (Fig. 6b). This suggests that suboptimal surgical procedures are not the exclusive source of targeting inaccuracy, but precision is further limited by the individual anatomical variability of the target coordinates, as well as deviations of the bone-defined Bregma point from the idealized origin of the atlas coordinate system^34–36^. Applying CT-MR fusions, one might adjust Bregma position or optimize target coordinates for each animal individually, as implemented in human surgical planning and in some monkey implantation surgeries^21,22,37,38^. Thus, pre-operative imaging for surgical planning can further save time and resources, especially in studies where animals have been pre-trained on a behavioral task before the surgery or the availability of subject animals is limited.

Since CT-based *in vivo* implant localization has not yet been performed in mice, we investigated the incurred radiation dose to provide recommendations for parameter settings that minimize radiation without compromising localization accuracy. We optimized our scanning procedures to provide sufficient quality without adverse effects on health including weight, integument or performance in behavioral tasks. We provided recommendations for acquisition settings for pre- and post-operative micro-CT scans based on the ALARA principle (‘as low as reasonably achievable’) for human radiation protection^39^ (Fig. 8).

Non-invasive, *in vivo* imaging modalities are essential to the longitudinal study of animal models in many biomedical research areas including oncology, neurology, cardiology and drug research^40–44^. We believe that the *in vivo* localization method presented here might become a similarly important tool in longitudinal neurophysiology experiments. With the widespread use of small animal imaging techniques in most fields of biomedical sciences^45,46^, preclinical CT and MRI equipment are readily accessible at many universities and research institutes. In addition, functional MRI measurements^47–50^ have already facilitated the installation of MRI machines, capable of high precision implant localization, at a large number of neuroscience research centers. Nevertheless, we showed that micro-CT imaging in itself might be sufficiently precise for part of the applications.

## Methods

### Animal care and use

Wild type (n = 24) and genetically modified (ChAT-Cre, n = 18; ChAT-Cre × DAT-Cre, n = 4) adult (3-4 months old) male mice of C57BL/6N genetical background were used. Animals were housed individually under a standard 12 hours light-dark cycle with food and water available *ad libitum*. All experiments were approved by the Committee for the Scientific Ethics of Animal Research of the National Food Chain Safety Office (PE/EA/675-4/2016, PE/EA/1212-5/2017, PE/EA/864-7/2019) and were performed according to the guidelines of the institutional ethical code and the Hungarian Act of Animal Care and Experimentation (1998; XXVIII, section 243/1998, renewed in 40/2013) in accordance with the European Directive 86/609/CEE and modified according to the Directives 2010/63/EU.

### Stereotaxic Surgery

Animals were anaesthetized using an intraperitoneal injection of 25 mg/kg xylazine and 125 mg/kg ketamine in 0.9% NaCl. The skin and subcutaneous tissues of the scalp were infused by Lidocain to achieve local anesthesia and eyes were protected with ophthalmic lubricant (Corneregel eyegel, Bausch & Lomb, Rochester, NY, USA). The mouse was placed in a stereotaxic frame (David Kopf Instruments, Tujunga, CA, USA) and the skull was levelled. The skin was incised, the skull was cleaned and a cranial window was opened above the target area (HDB, +0.74 mm antero-posterior, 0.6 mm lateral; MS, +0.9 mm antero-posterior, 0.9 mm lateral; VTA; −3.1 mm antero-posterior, 0.6 mm lateral; ventral CA3 −2.5 mm antero-posterior, 2 mm lateral). Injections of the adeno-associated virus vector encoding ChR2 (AAV2.5.EF1a.Dio.hChR2(H134R)eYFP.WPRE.hGH, 300 nl, titer: 7,7*10^12^ GC/ml); Penn Vector Core, PA, United States) were performed using glass pipettes pulled from borosilicate capillaries broke to 20–30 μm tip diameter, connected to a MicroSyringe Pump Controller (World Precision Instruments, Sarasota, FL, USA), lowered stereotaxically into the target nucleus (HDB, DV −5.0/−4.7 mm; MS, DV, −3.5/−4.2/−4.6 mm; VTA, DV −4.4/−4.0 mm, while in case of the CA3 no virus injection was performed).

Custom-built microdrives (see Hangya et al.^33^ for details on fabrication) housing 8 nichrome tetrodes (diameter, 12.7 μm; Sandvik, Sandviken, Sweden) and a 50 μm core optic fiber (outer diameter, 65 ± 2 μm; Laser Components GmbH, Olching, Germany); 105 μm core optic fibers (outer diameter, 250 ± 4 μm; Thorlabs Corp., Munich, Germany); or a silicon probe (Buzsáki type, Neuronexus Technologies A1×16, Ann Arbor, MI, USA, with a single 52 × 15 μm shank) were implanted in (electrode drives) or above (fiber optics) the target nuclei without using a guiding cannula, as described previously^33^.

Fluorescent dye (DiI; Invitrogen LSV22885, Ottawa, Canada) was applied on the tip of the tetrode implants before lowering for later histological localization. Implants were secured by acrylic resin (Jet Denture; Lang Dental, Wheeling, USA). Buprenorphine was used for post-operative analgesia (Bupaq, 0.3mg/ml; Richter Pharma AG, Wels, Austria).

### Behavioral training

Mice were trained on the 5-choice serial reaction time task described by Bari et al.^31^. Instead of food reward, mice were water restricted to approximately 85-90% body weight and received water reward from the same port where the cue was presented during the training. All mice were pre-trained before the surgery in a custom-built automated training system where they were allowed to perform the task several times per day, until they reached 80% performance (5-10 days). After one week of recovery period following the surgery, post-operative CT scans were performed in case of the test group, and the behavioral training was resumed.

### Atlases

The Bregma and Lambda points and the axes of the stereotaxic coordinate system were defined according to Paxinos and Franklin^27^. Bregma and Lambda were determined as ‘the midpoints of the curve of best fit along the coronal and the lambdoid suture, respectively. They are not necessarily the points of intersection of these sutures with the midline suture’. The same atlas was used for surgical planning and comparing coordinates across localization methods. For localization based on CT-MRI fusion, a three-dimensional mouse brain atlas by Bai and colleges^28^ was used, which is based on the T2-weighted MRI images of 5 mice with structures manually identified on the basis of the Paxinos atlas.

### Pre-operative scanning

Pre-operative CT and MRI scanning was performed in the same scanning bed to avoid postural changes and disparities between the imaging planes. Isoflurane gas was used as an inhalation anesthetic through a specialized isoflurane mask which was also used for head fixation to avoid motion artifacts.

CT measurements were performed on a NanoX-CT (Mediso, Budapest, Hungary) cone-beam micro-CT imaging system with an 8W power X-ray source. Several acquisition parameters were tested for image quality and radiation dose (Fig. 8a). We used simple circular scanning to scan the head of the animal only, to further minimize radiation dose. Pre-operative CT imaging parameters were set to minimize the radiation dose, since all tested settings provided sufficient quality images for localizing bone landmarks: tube voltage, 45 kVp; magnification, 1.36; exposure time, 500 ms; number of projections, 180. These represent minimal values within the adjustable range with 3-minute acquisition time.

Magnetic resonance imaging was performed with a nanoScan PET/MRI system (Mediso, Budapest, Hungary), which is equipped with a permanent magnetic field of 1T and with a 450 mT/m gradient system using a volume coil for both reception and transmission. T_1_-weighted 3D gradient echo sequence was acquired using 8 excitations, T_R_=15 ms repetition and T_E_=2.2 ms echo times and 25° flip angle with the resolution set to 0.28 mm. The sequence parameters were selected in order to achieve proper contrast and SNR that facilitates good visualization of the contour of the brain and the position of the ventricles with sufficient resolution in reasonable acquisition time (10 minutes).

### Post-operative scanning

Post-operative CT scans were performed using a similar procedure as detailed above 4-12 days after the surgery. In case the implant was lowered during the *in vivo* electrophysiology recordings, the post-operative CT scanning was repeated after the termination of the experiments for accurate comparison of *in vivo* localization with the histology results (see ‘Quantification of localization accuracy using gold-standard histology’ section). Metal parts of the drive were positioned to be pointing away from the plane in which the X-ray source is rotating, so their shadow-artifacts bypassed our region of interest, i.e. the close environment of the implant. To examine the image quality - radiation dose trade-off several post-operative acquisition settings were tested in the range of 500-1300 ms exposure time; 180-360 projections; 1.36-2.47 magnification, with the tube voltage set to 45 kVp. Optimal acquisition settings depending on the size and radiodensity of the implant are provided in the ‘Optimization of image quality and radiation dose’ section of Results. Due to the presence of the metal parts, no MRI scanning was performed after the surgery.

### Image reconstruction and registration

For CT image reconstruction, we used filtered back projection with a Butterworth filter^51^, and the isotropic voxel size was set to the minimum value of 35 μm or 19 μm for 1.36 and 2.47 magnifications, respectively. Coordinate system transformations and the CT-MRI-atlas co-registrations were performed with the VivoQuant (inviCRO, Boston, MA, USA) pre-clinical medical image post-processing software, using Euclidean transformations (translation and rotations, except for the step (ii) of the CT-MRI fusion method where non-Euclidean affine transformations were used), based on anatomical landmarks. Electrode tracks were segmented with an intensity threshold applied on a whole-brain ROI.

### Histological track reconstruction

After termination of the experiments, mice were deeply anesthetized with ketamine-xylazine (see above). In animals implanted with tetrode drives, to accurately mark the position of the electrode tip for histological reconstruction, electrical lesioning (40 μA for 5 seconds applied through 1 or 2 leads; IonFlow Bipolar, Supertech Instruments, Pécs, Hungary) was performed. Mice were then perfused transcardially with 0.1M phosphate buffered saline (PBS) for 1 min, then with 4% paraformaldehyde (PFA) in PBS for 20 min. Implants were carefully removed, brains were extracted from the skull and post-fixed for 24 h in PFA at +4°C.

50-μm-thick coronal sections were cut (Vibratome VT1200S, Leica, Wetzlar, Germany), mounted on microscope slides, covered in mounting medium (Vectashield, Vector Labs) and examined with a fluorescent microscope (Nikon Eclipse Ni microscope). Darkfield, brightfield and fluorescent images were taken with a Nikon DS-Fi3 camera. Images were processed for full track reconstruction as described by Hangya et al.^33^. Briefly, mouse brain atlas sections^27^ were co-registered with the corresponding coronal slices using linear scaling and rotation based on anatomical landmarks identified on the bright field and dark field images. The atlas images were carried over to fluoromicrographs that visualized the infected area by virally transfected ChR2-eYFP and the trajectory of the implant by fluorescent red Dil applied on the implants. Implant tip coordinates and trajectory directions were determined based on the Dil marked tracks and the electrical lesions localized in the bright field and dark field images.

### Radiation dose measurements

Calibrated lithium fluoride MCP-N (LiF:Mg,Cu,P) thermoluminescent dosimeters (TLD) were used to measure radiation exposure. These type of TLD chips have a broad linear dose range^52^, from cGy-s to approximately 10 Gy. Before going through the CT scanning, TLD detectors were attached to the neck of the animals as close to the skull as possible. After the irradiation, radiation doses were read from the detectors with a TLD Cube (RadPro, Wermelskirchen, Germany) reader system.

### Statistics

Correlations were tested using Pearson’s correlation coefficient (r), at a significance level of 0.05. The error measures were tested for statistical significance by two-tailed Wilcoxon signed rank test. Systematical deviations from the target coordinate and directions were tested with one-tailed *t*-test. Statistical analyses were carried out in Matlab (The Mathworks, Natick, MA, USA).

## Data availability

The data that support the findings of this study are available within the article and the associated Supplementary material. Any other data are available from the corresponding author upon reasonable request.

## Acknowledgements

We thank Gabriella Taba for the TLD detectors and help with the radiation dose measurements; Dr. Andor Domonkos for providing the silicone probe; Drs. Sergio Martínez-Bellver, Viktor Varga and Duda Kvitsiani for helpful comments and discussions on the manuscript. This work was supported by the ‘Lendület’ Program of the Hungarian Academy of Sciences (LP2015-2/2015), NKFIH KH125294 and the European Research Council Starting Grant no. 715043 to BH and by the ÚNKP-19-3 New National Excellence Program of the Ministry for Innovation and Technology to BK. BH is a member of the FENS-Kavli Network of Excellence.

## Author information

### Affiliations

Lendület Laboratory of Systems Neuroscience, Institute of Experimental Medicine, Hungarian Academy of Sciences, Szigony u. 43. H-1083 Budapest, Hungary

Bálint Király, Diána Balázsfi, Nicola Solari, Katalin Sviatkó, Katalin Lengyel, Eszter Birtalan & Balázs Hangya

Department of Biophysics and Radiation Biology, Semmelweis University, Tűzoltó u. 37-43, H-1094 Budapest, Hungary

Bálint Király, Ildikó Horváth, Domokos Máthé & Krisztián Szigeti

Department of Biological Physics, Eötvös University, Pázmány P. stny. 1A, H-1117, Budapest, Hungary Bálint Király

János Szentágothai Doctoral School of Neurosciences, Semmelweis University, Üllői út 26., H-1085, Budapest, Hungary

Katalin Sviatkó

CROmed Translational Research Centers, Tűzoltó u. 37-43, H-1094, Budapest, Hungary Domokos Máthé

### Contributions

The idea was developed by BH and BK. BK and IH designed the scanning protocol and performed the *in vivo* imaging experiments. BK developed the *in vivo* localization protocol, performed the radiation dose measurements, analyzed the data and prepared the figures; DB, NS, KS and BK performed the implantation surgeries; DB, KL and BK performed histological localization; DB and EB performed the behavioral tests. The project was supervised by BH, KSz and DM. The manuscript was written by BK, BH and DB with comments from all authors.

### Corresponding author

Correspondence to Balázs Hangya

